# Loss of altruism in the social amoeba *Dictyostelium discoideum* is associated with the G protein-coupled receptor *grlG*

**DOI:** 10.1101/2022.10.21.513250

**Authors:** Laura M. Walker, Rintsen N. Sherpa, Sindhuri Ivaturi, Debra A. Brock, Jason R. Walker, Joan E. Strassmann, David C. Queller

## Abstract

Aggregative multicellularity relies on cooperation among individual cells to form a multicellular body. In *Dictyostelium discoideum* this cooperation is maintained by high relatedness. Previous work showed that experimental evolution under low-relatedness resulted in an increase of cheaters (cells that contribute proportionally more to spores than to the sterile stalk) and that many clones completely lost cooperation and the ability to form fruiting bodies. Here, we investigate the genomic changes underlying the evolution of the cheating phenotype using whole-genome sequencing and variant analysis of these previously evolved *D. discoideum* lines. We identified 38 single nucleotide polymorphisms in 29 genes, none of which have been previously implicated in cheating. Each gene has one variant except for the G protein-coupled receptor *grlG*, which has at least one variant in over half of the lines. Upon identifying the parallel evolution of *grlG*, we screened additional clones to investigate the correlation between variants in the gene and the loss of cooperation (identified by the inability to form a fruiting body). We found that variants in the 5’ half of *grlG* that impact the signal peptide or extracellular binding domain are significantly associated with the loss of cooperation (non-fruiting); the association was not significant in the 3’ half of the gene. This suggests that the loss of *grlG* was adaptive under low-relatedness and that the 5’ half of the gene in particular is important for cooperation and multicellular development. This confirms the importance of high relatedness in the evolution of altruism in the social amoeba *D. discoideum*.

## Introduction

High relatedness is crucial for the maintenance of cooperation in multicellularity. Without high relatedness to reduce conflict among individuals in a cooperative group, cheaters can evolve that exploit the individual and destabilize altruistic traits (Hardin 1968; Gilbert *et al*. 2007). High relatedness is easily achieved within organisms that go through a single-cell bottleneck or are otherwise clonal, as is true for most of the more than 20 transitions from unicellularity to multicellularity (Grosberg and Strathmann 2007; Knoll 2011). A single-cell origin ensures that all cells are highly related (genetically identical) which reduces conflict, protects against cheaters and promotes the division of labor that is so crucial for complex multicellularity such as in plants and animals (Grosberg and Strathmann 2007; Cooper and West 2018). However, a single-cell origin is not the only way to achieve multicellularity and its many benefits. Many other organisms from across the tree of life achieve multicellularity via aggregation or fusion of individual cells (Bonner 1998; Sebé-Pedrós *et al*. 2017). In this form of multicellularity, called aggregative multicellularity, the cells that aggregate need not be related, so other mechanisms are required to reduce conflict and maintain cooperation (Queller 2000).

Organisms that arise by aggregation offer important opportunities for evolutionary and mechanistic studies of conflict and cooperation. Genetic indications of conflict and its control are likely to be evident in multicellular organisms that arise by aggregation. Unlike multicellular organisms that require a single-cell bottleneck, in organisms with aggregative multicellularity it is possible to experimentally manipulate intra-organismal relatedness and study the outcome. Among the best studied organisms with aggregative multicellularity is the social amoeba *Dictyostelium discoideum.* Usually free-living in the soil, when faced with starvation *D. discoideum* amoebae aggregate to cooperatively form a multicellular fruiting body in which about 80% develop into reproductive spores and the remaining 20% die forming a stalk (Kessin 2001). The stalk lifts the spores, facilitating dispersal (smith *et al*. 2014). Stalk formation is altruistic because cells that contribute to the stalk die, sacrificing themselves to aid in spore dispersal. This partitioning of reproductive spores and sterile stalk cells provides an opportunity for clones to cheat in mixtures of genotypes by contributing more than their fair share of cells to spores versus the stalk (skewing the typical distribution) (Strassmann *et al*. 2000).

*Dictyostelium discoideum* is a particularly valuable and tractable model system for studying aggregative multicellularity (Strassmann and Queller 2011; Ostrowski 2019). In the laboratory we can experimentally manipulate populations of genetically distinct amoebae and they will aggregate together and develop into a fruiting body. An extensive set of experimental tools and resources (fluorescent labels, reference genome, etc.) are available that allow us to track the outcome of the distinct genotypes (Buttery *et al*. 2013). In the multicellular fruiting bodies that result from mixing experiments, it is usually possible to identify a winner (the clone contributing more to spores than stalk) and a loser (the clone contributing more to the stalk than spores) (Strassmann *et al*. 2000; Fortunato *et al*. 2003a; Buttery *et al*. 2009; Wolf *et al*. 2015; Madgwick *et al*. 2018). Evidence suggests that cheating occurs in natural populations too. Despite the high relatedness in natural populations (Gilbert *et al*. 2007), distinct clones are often found in close proximity and they too will aggregate and form chimeric fruiting bodies (Strassmann *et al*. 2000; Fortunato *et al*. 2003b; smith *et al*. 2016).

Genetic mutations that result in a selfish trait such as the preferential formation of spores versus stalk (cheater mutations), may pose a threat to cooperation depending on the context in which they arise. Within an individual fruiting body a cheater mutation can be favored by selection and contribute disproportionately more to spores than to stalk (Strassmann *et al*. 2000). But the outcome more broadly will depend on the relatedness of aggregating cells and the extent to which those carrying the mutation can still cooperate on their own. If a clone carrying the cheater mutation has completely lost cooperation such that it can only form spores (an obligate cheater), then it can only persist when an exploitable clone is available. When an obligate cheater clone finds itself alone without others to form a stalk, multicellular fruiting body formation (development) comes to a halt resulting in a mound of cells that cannot disperse, and thus has a fitness of zero. We call these obligate cheater clones “non-fruiters.” If instead, a clone carrying the cheater mutation can still cooperate and can modify its strategy depending on the availability of exploitable clones then the mutation may persist without threatening the collapse of multicellularity. We call these clones “facultative cheaters.”

It has been experimentally shown that selection under low relatedness increases the occurrence of both facultative (Santorelli *et al*. 2008) and non-fruiting obligate cheaters (Ennis *et al*. 2000; Kuzdzal-Fick *et al*. 2011). When obligate cheaters arise in the wild where relatedness is high, they should not persist because they will usually find themselves in clonal or nearly clonal aggregations where their cheating advantage will not trump the disadvantage of their inability to form fruiting bodies on their own (Fortunato *et al*. 2003b; Gilbert *et al*. 2007). In line with that expectation, no obligate cheaters have been identified from natural populations of *D. discoideum*. While it is true that non-fruiting clones would be missed with traditional isolation techniques (Cavender and Raper 1965), Gilbert *et al*. (2007) screened specifically for the presence of non-fruiting obligate cheaters by clonally plating over 3,000 spores from wild-collected fruiting bodies to observe the phenotype. No clones were found that were incapable of forming a fruiting body. These findings are in line with predictions of kin selection theory (Hamilton 1964) and show that high relatedness helps stabilize altruism in *D. discoideum.* The altruistic trait of stalk formation can be maintained only if the benefits are shared with related individuals who likely share the altruism allele.

One way in which organisms with aggregative origins of multicellularity maintain high relatedness and the resulting cooperation is through kin discrimination (West *et al*. 2006). Kin discrimination describes mechanisms that allow individuals to identify and preferentially direct benefits to relatives and either harm or ignore non-relatives (Tsutsui 2004; Strassmann *et al*. 2011). Some species of Dictyostelids have robust kin discrimination systems that aid in the sorting of different genotypes into mostly clonal fruiting bodies (Mehdiabadi *et al*. 2006, 2009; Sathe *et al*. 2014). *Dictyostelium discoideum* also has kin discrimination with sorting regulated by two highly variable cell adhesion genes, *tgrB1* and *tgrC1* (Benabentos *et al*. 2009; Hirose *et al*. 2011; Gruenheit *et al*. 2017). Segregation by kin discrimination in *D. discoideum* takes place during early aggregation but is not maintained and does not result in fruiting bodies that are highly sorted by relatedness (Flowers *et al*. 2010; Gilbert *et al*. 2012). So although it may limit some cheaters from invading (Ho *et al*. 2013), kin discrimination alone is not strong enough to provide complete protection from selfish cheaters in *D. discoideum*. Instead, high relatedness in natural populations must be maintained in additional ways such as through structured population growth and spatial distribution patterns (Buttery *et al*. 2012; smith *et al*. 2016).

Despite the great deal has been learned about cooperation and conflict in *D. discoideum,* much remains to be learned with respect to the underlying genes and pathways. Several efforts to identify genes underlying cheating phenotypes have been made, often by employing restriction enzyme-mediated integration (REMI) to generate gene knock-out mutant libraries (Kuspa and Loomis 1992). Screening of such REMI mutant libraries has successfully revealed a number of genes involved in cheating phenotypes in *D. discoideum.* Santorelli *et al*. (2008) screened a large REMI mutant library, focusing on facultative cheater genes (by requiring one round of clonal fruiting body development) and identified over 100 genes predicted to cause cheating when lost to mutation. As might be expected for such a large number of loci, the gene ontology terms span a wide diversity of biological processes. Thus far only two of the genes from Santorelli *et al*. (2008) have been characterized including *chtB* (Santorelli *et al*. 2013) and *chtC* (Khare and Shaulsky 2010). Despite the expectation that mutations in genes that cause cheating should have an associated fitness cost to control their spread, neither the *chtB* nor *chtC* mutants have any obvious fitness costs. Mutant clones for both genes are able to cheat facultatively, increasing in numbers in chimera but when alone, they contribute cells to both spores and stalk and form fruiting bodies normally (Khare and Shaulsky 2010; Santorelli *et al*. 2013). It may be that there are disadvantageous pleiotropic effects that keep these mutations from spreading, but they were not detected in the laboratory. Pleiotropy is, after all, another mechanism proposed to hinder the spread of cheaters such as described for two separately identified facultative cheater genes in *D. discoideum*, *dimA* and *csaA* (Queller *et al*. 2003; Foster *et al*. 2004; Strassmann and Queller 2011).

In addition to the aforementioned facultative cheater genes, only one obligate cheater gene has been described, *fbxA* (Ennis *et al*. 2000). The *fbxA* mutant, also resulting from a REMI mutant library screen, was the first cheater gene to be described and has therefore also been referred to as “Cheater A” or “chtA” for short. When cells from a clone carrying a mutant copy of *fbxA* (encoding F-box protein A) are mixed with wild-type cells, they become overrepresented as spores rather than stalk cells (Ennis *et al*. 2003). But unlike the facultative cheaters, there is a large group cost; as the frequency of *fbxA* mutants in a group increases, the overall spore production decreases (Gilbert *et al*. 2007). And when alone (*i.e.,* only *fbxA* mutants are present), few to no spores are produced; the loss of *fbxA* results in non-fruiting obligate cheaters (Ennis *et al*. 2000, 2003).

The cheater genes that have been described thus far in *D. discoideum* share few sequence features or protein domains and are associated with a wide diversity of cellular functions and pathways, which suggests that cheating can be accomplished in many ways (Santorelli *et al*. 2008). Some of the mechanistic diversity is already apparent; the described cheater genes cause cheating via different means including altered communications in cell-fate determination (Ennis *et al*. 2000; Thompson *et al*. 2004; Khare and Shaulsky 2010) and changes in cell adhesion (Queller *et al*. 2003). Perhaps that is to be expected given the complexity of multicellular development in *D. discoideum*. In addition to the aggregation and cooperation required of the previously free-living individuals, multicellular development involves the differentiation of cell types (*e.g.,* prestalk, prespore), followed by the maintenance and coordination of each of those cell types fates and their proportions (Loomis 2015). Cheaters can presumably arise that exploit any number of aspects of this developmental process; the study of cheating therefore also has the potential to provide insight into development more broadly.

To expand our understanding of the genomics underlying cheating and cooperation in *D. discoideum,* and especially the understudied obligate cheaters, in this study we return to cell lines experimentally evolved under low-relatedness by Kuzdzal-Fick *et al*. (2011). Starting from a single isolate of the wild type lab strain AX4, Kuzdzal-Fick *et al*. (2011) established 24 replicate lines and maintained low-relatedness by transferring a million spores to new plates 31 times (about 290 generations) (Figure 1A). Each round began with one million spores which were allowed to hatch, proliferate by eating bacteria, and then form fruiting bodies. At each transfer, the low relatedness was reestablished by replating a million thoroughly mixed spores. In this way, cheaters that appeared by mutation would not be with other such cells among the million and would instead be in close proximity to cells to exploit. Under these conditions, cheaters were favored by selection and significantly increased in abundance by contributing more to spores while contributing less to the stalk. As the percentage of cheaters in a line increased, fewer spores were produced overall. And among the evolved lines, the percentage of clonal isolates which had completely lost cooperation and could no longer form fruiting bodies on their own (non-fruiters) rose to as high as 69% (Figure 2). On average, the evolved lines of Kuzdzal-Fick *et al*. (2011) contained approximately 30% non-fruiters and most of them that have been tested were cheaters.

**Figure 1.**
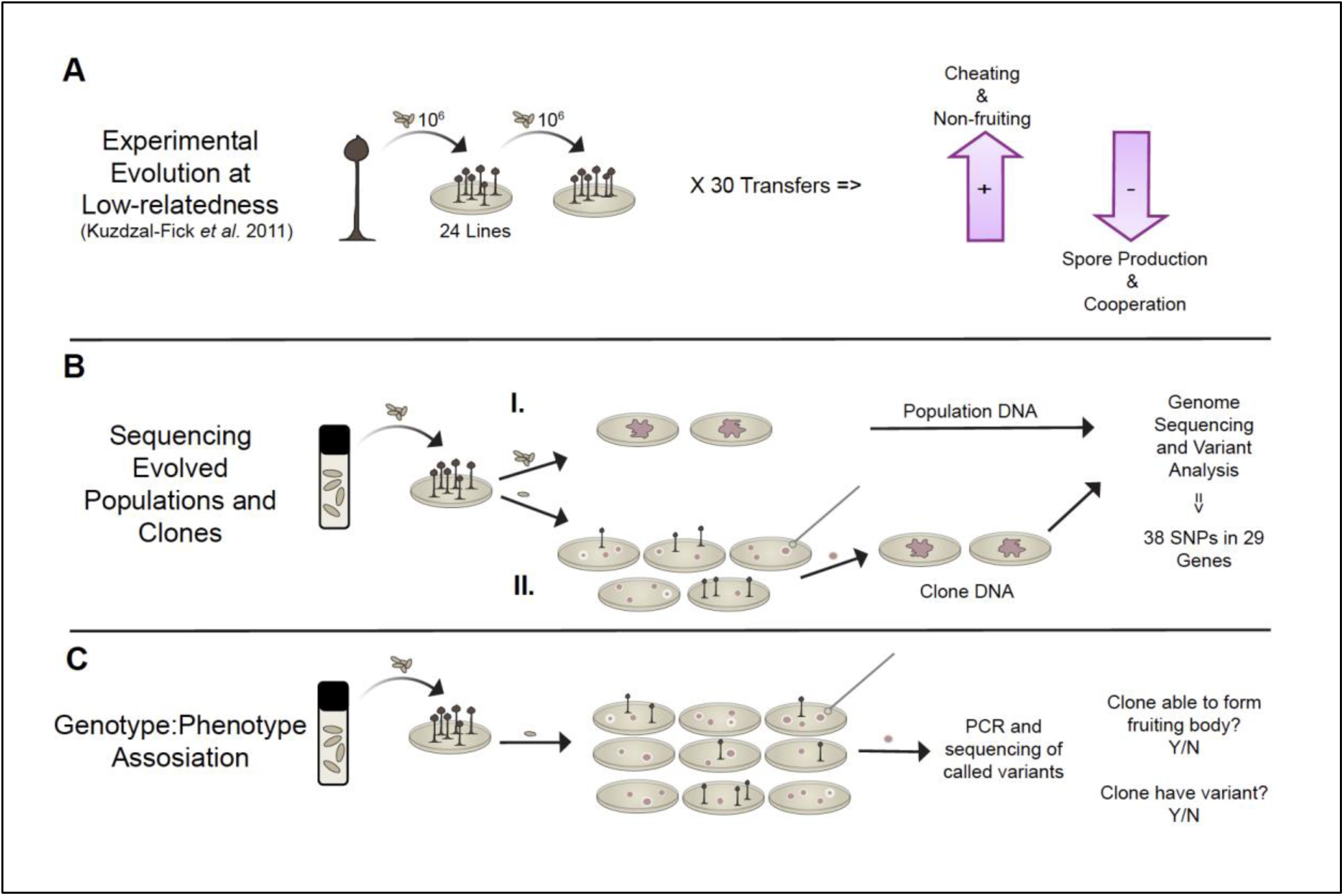
Outline of the experimental workflow. **A.** Experimental evolution of *D. discoideum* at low-relatedness by Kuzdzal-Fick *et al*. (2011). A clonal isolate of AX4 was used to generate 24 replicate lines. After fruiting body formation, spores were collected, thoroughly mixed and one million spores were replated. Reestablishing low relatedness at each of the 31 passages (about 290 generations) allowed selection to favor mutations that conferred cheating. **B.** Whole-genome sequencing and variant analysis in this study of the 24 evolved cell lines (from A). We grew spores from the final passage of each evolved cell line to generate two samples, (**I.**) the line in bulk, as a population and (**II.**) a non-fruiting clone from each line. For the line in bulk as a population (**B, I.**), we grew cells for genomic DNA extraction. For the non-fruiting clone (**B, II.**), we plated a serial dilution of spores from each population to isolate a non-fruiting clone which we then grew for genomic DNA extraction. We used whole-genome sequencing and variant analysis of populations and clones to identify 38 SNPs impacting 29 genes. **C.** Association of SNPs with the loss of altruism in clones. We again plated the evolved populations (lines) (from A) to allow growth of clonal colonies from individual spores. We evaluated numerous clones from each line by two criteria: 1) whether they were obligate cheaters (non-fruiters) or not (normal fruiting body formation) and 2) whether they carried the previously identified SNP or not, via PCR and Sanger sequencing.

**Figure 2.**
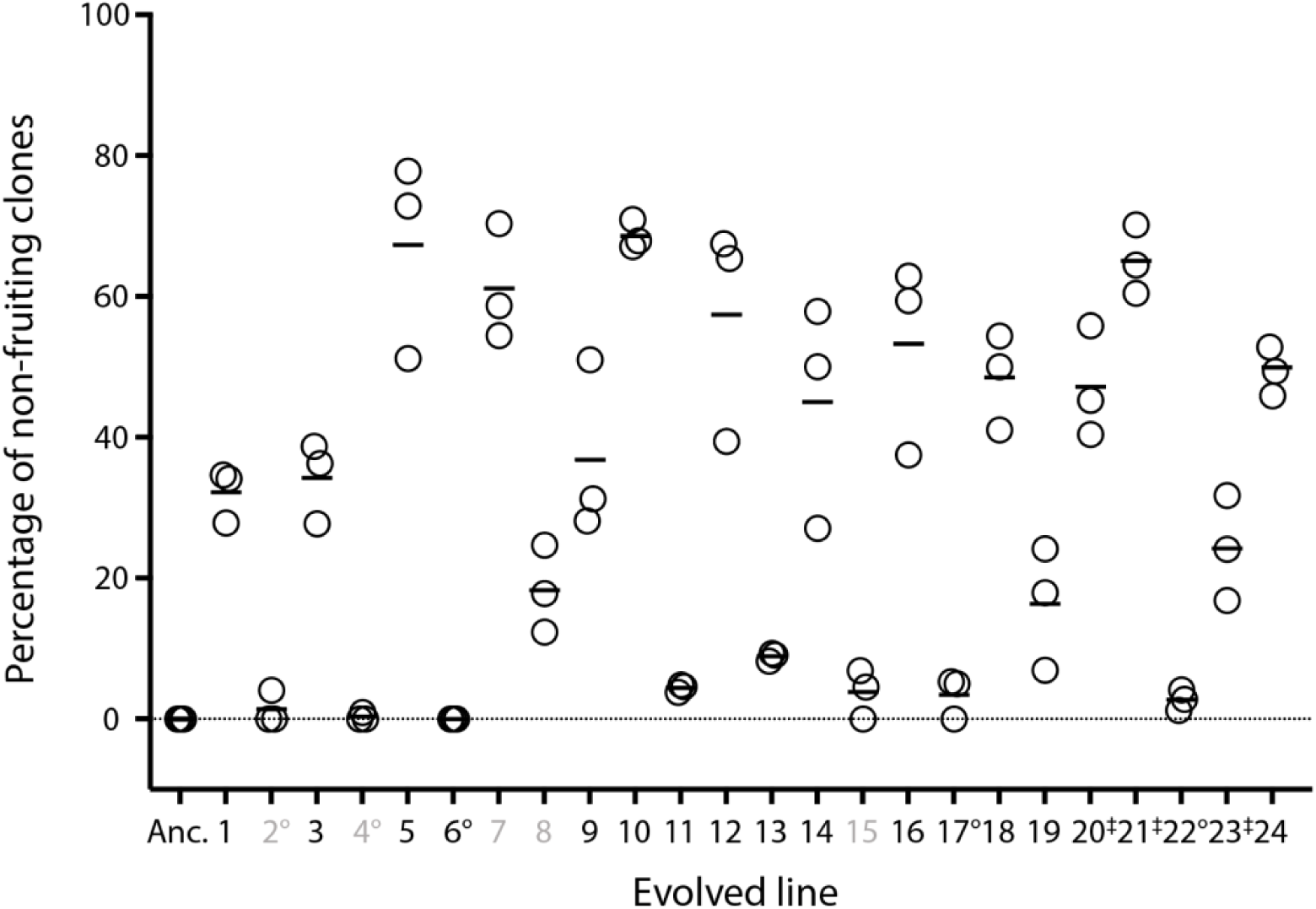
Percentage of non-fruiting clones per evolved cell line from Kuzdzal-Fick *et al*. (2011). After about 290 generations under low-relatedness, 19 of the 24 lines had evolved to cheat their ancestor (the five lines that did not cheat are labeled in grey). Non-fruiting (obligate cheating), which was not present in the ancestor (Anc.), also increased in frequency to a varying degree among the evolved lines. The percentage of non-fruiting clones for each line (three replicate measurements) is displayed in this scattered dot plot with the mean represented by the horizontal bar. In this study we generated a whole-genome sequence for each evolved line in bulk as a population and for one non-fruiting clone except where indicated (’°’ indicates no clone was sequenced and ‘‡’ indicates two clones sequenced). Redrawn from Kuzdzal-Fick *et al*. (2011).

The work by Kuzdzal-Fick *et al*. (2011) showed that drastically reduced relatedness allowed the spread of cheater mutations that contribute disproportionately to spores and reduced the overall spore production in many lines (Figure 1A). This demonstrates how low relatedness in a natural population would lead to a collapse of multicellularity and with it, the advantages of fruiting body formation and spore dispersal. In this study we use whole-genome sequencing and variant analysis of these experimentally evolved *D. discoideum* cell lines to identify genomic changes that allowed non-fruiting obligate cheaters to evolve over the course of that experiment.

## Materials and Methods

### Experimental Cell Lines

*Dictyostelium discoideum* cell lines were experimentally evolved under conditions of low-relatedness by Kuzdzal-Fick *et al*. (2011) (Figure 1A). Kuzdzal-Fick *et al*. (2011) froze spores from the evolved lines in KK2 buffer (2.25 g KH_2_HPO_4_ and 0.67 g K_2_HPO_4_ per L) with 25% glycerol and stored them at −80°. We thawed spores from the ancestor and from the final passage of each evolved line for genomic DNA extraction. Each evolved line is a population composed of cells and lineages carrying any newly acquired mutations. To capture all of this variation, we generated a whole-genome sequence for each evolved line. We will refer to these sequences as populations. To narrow the focus to variation associated with obligate cheating, we also generated a whole-genome sequence for one non-fruiting clone from each evolved line (Figure 1B). We will refer to these sequences as non-fruiter clones. And finally, to identify variation that arose during the experimental evolution we generated a whole-genome sequence of the ancestor.

To isolate genomic DNA from each of the 24 evolved lines (Figure 1B, I.), we plated spores onto two SM/5 agar plates (2 g glucose, 2 g BactoPeptone (Oxoid), 2 g yeast extract (Oxoid), 0.2 g MgCl_2_, 1.9 g KH_2_PO_4_, 1 g K_2_HPO_4_ and 15 g agar per liter) with 200 µL of *Klebsiella pneumoniae* in KK2 buffer (OD_600_ 1.5) as food. We plated the spores at two different concentrations (2 × 10^5^ spores and 4 × 10^5^ spores per plate) to allow processing of multiple samples at one time in case growth rates varied between the evolved lines. We incubated the plates at room temperature for approximately 36 h or until log phase. We collected amoebae in log phase from the surface of two plates for each sample and washed them four to five times in chilled KK2 buffer to remove the food bacteria before DNA extraction.

To isolate genomic DNA from non-fruiting clones (Figure 1B, II.) we first plated serial dilutions of spores to allow for clonal growth from individual spores. We aimed to collect one non-fruiting clone from each evolved line but for five lines (2, 4, 6 and 17 and 22), despite repeated attempts, we were unable to locate any non-fruiting clones. To use the remaining sequencing space, we sequenced a second non-fruiting clone for three haphazardly selected lines (20, 21 and 23). After three to five days of growth, we collected cells from the leading edge of each non-fruiting clonal plaque using a sterile loop and plated them on SM/5 agar with *K. pneumoniae*. We then plated these cells to grow for DNA extraction using the same protocol described for the evolved lines. To document the non-fruiting morphology, we took photographs of each clonal plaque immediately before collecting cells from the leading edge for expansion (example images are available in Supplemental Material 1). A second photograph was taken one to two days following collection and we continued to monitor the plaques to ensure they never formed fruiting bodies.

### DNA isolation and sequencing

We isolated genomic DNA from the washed, log-phase cells in the range of 1–2 × 10^8^ cells using the Qiagen DNeasy® Blood and Tissue kit (Qiagen). We resuspended the genomic DNA in 10 mM Tris-HCl pH 8.5 and stored it at 4° until submission to the McDonnell Genome Institute at Washington University in St. Louis, MO for library preparation and sequencing. Sequence libraries were prepared starting with 0.5 μg of genomic DNA using the KAPA Hyper Library Prep (KAPA Biosystems). We sequenced genomic DNA on the Illumina NovaSeq (150 bp x 2 paired-end) to an estimated depth of 100X and 500X for the clones and populations, respectively.

### Sequence Alignment

We aligned the Illumina paired-end reads to a single FASTA file containing the reference genomes of both *D. discoideum* AX4 and the food bacterium, *K. pneumoniae* (both downloaded from NCBI in June 2019). We used BWA-MEM (0.7.15) (Li 2013) to index the *D. discoideum* and *K. pneumoniae* concatenated reference genome and to align the paired-end reads for each sample. We used an alignment pipeline to run BWA-MEM (Li and Durbin 2009) which converts, sorts and indexes input, intermediate, and output formats using Picard v2.18.1 (http://broadinstitute.github.io/picard/), Sambamba v0.6.4 (Tarasov *et al*. 2015) and Samtools v1.3.1 (using HTSLib v1.3.2) (Li *et al*. 2009) ultimately resulting in an aligned, sorted, compressed and indexed CRAM file. We used both Picard (CollectInsertSizeMetrics, CollectAlignmentSummaryMetrics, CollectGcBiasmetrics) and Samtools (flagstat) to evaluate alignment and coverage metrics using BAMs of the initial alignments including both *D. discoideum* and *K. pneumoniae* reference genome alignments and again after excluding reads that aligned to *K. pneumoniae*.

### Variant Calling and Filtration with GATK

We calculated the initial genotype likelihoods using GATK (4.1.2) (McKenna *et al*. 2010) HaplotypeCaller (-ERC GVCF --sample-ploidy 1) for each sample. Next, we ran GATK GenotypeGVCFs (--sample-ploidy 1) to create per-sample genotypes as individual VCF files for the entire *D. discoideum* and *K. pneumoniae* concatenated reference genome. We selected the *D. discoideum* chromosomes (and the unplaced contigs associated with the reference genome) from the VCF files for downstream annotation and filtering using GATK SelectVariants. We used the Ensembl Variant Effect Predictor (VEP 95.3) (McLaren *et al*. 2016) to annotate all variants and add sequence ontology terms (--term SO) using the dicty2.7 assembly of ‘*Dictyostelium*_*discoideum*’ from Ensembl Protists (release 43). We decomposed complex variants using vt decompose (Tan *et al*. 2015) before adding allele frequencies, merging or filtering. We then processed the decomposed variants with bam-readcount (0.7.4) (https://github.com/genome/bam-readcount) and cyvcf2 (https://github.com/brentp/cyvcf2) to generate allele frequencies. We used VAtools (https://github.com/griffithlab/VAtools, 3.1.0) to add the AF (allele frequency) format fields to each sample including the allele frequencies of each allele as calculated by bam-readcount allele counts. We processed all per-sample VCFs with bgzip and tabix for speed and storage before merging to generate the full per-sample call set VCF using GATK (3.6) CombineVariants (-genotypeMergeOptions UNIQUIFY).

We performed variant filtration of the GATK VCF using bcftools v1.12 (using HTSLib v1.12) (Li 2011) and GATK. We did not consider indels or sites with more than one alternate allele. We are only interested in variation that arose during the course of experimental evolution or, new variation between the ancestor and the evolved lines. To exclude pre-existing variation we first removed sites (from all samples) for which the ancestor was called as a variant (*i.e.,* sites in the ancestor that differed from the reference genome) as well as sites that were left uncalled in the ancestor. To remove ancestral polymorphism we next calculated the major allele frequency (MAF) of all sites in the ancestor BAM file using bam-readcount (0.7.4) (https://github.com/genome/bam-readcount). Using that information, we then removed sites (from all samples) for which in the ancestor, the MAF < 0.90.

We applied hard filters using GATK VariantFiltration and SelectVariants following the GATK Best Practices standard recommendations (QD > 2, FS < 60, SOR < 3, MQ > 40, MQRankSum > −12.5 and ReadPosRankSum > −8). Next we applied custom filters using bcftools view. The first custom filter was the removal of sites missing too much data, which we defined as sites left uncalled in more than 10 samples. For each of the 24 evolved lines we sequenced the whole line (as a population), and for most of the lines we also sequenced one non-fruiting clone, for a total of two samples per line (or a total of three samples for lines 20, 21 and 23 for which we sequenced two clones). There is a low likelihood that the same SNP will occur by chance in more than one evolved line. But because we sequenced two or three samples for each line (the population and individual clones) the maximum number of times that a SNP is likely to appear in our data is twice (or three times for lines 20, 21 and 23). For this reason, we applied a maximum alternate allele count (AC) of three. Next, we applied a minimum Phred-based quality score (QUAL) of 200 to remove low-quality sites and we removed sites with more than 1.5 times the average approximate read depth (DP) to reduce false positives. Last, we manually reviewed this final set of SNPs using the Integrative Genomics Viewer (IGV) to further reduce the number of false positives and misclassifications (Robinson *et al*. 2011, 2017).

### Variant Calling and Filtration with Freebayes

We also performed variant calling on all samples (joint calling) using Freebayes v1.3.1-dirty (Garrison and Marth 2012). We used the default parameters but for the following exceptions: sample ploidy of one, pooled continuous mode, minimum base quality 10, minimum mapping quality 10 and we only retained the best of six alleles. We streamed variant calls directly through the vcflib (Garrison *et al*. 2021) vcffilter, to remove variants with a quality score below 20. We decomposed complex variants into their constituent SNPs and indels using vcflib vcfallelicprimitives followed by normalization with vt (Tan *et al*. 2015).

We performed variant filtration of the normalized Freebayes VCF using bcftools v1.12 (using HTSLib v1.12) (Li 2011) following a similar process to that used for filtering the GATK VCF. Although the two callers calculate and output some different metrics for quality assessment, we generated filters resembling those applied to the GATK VCF as much as possible. First, we excluded indels and sites with more than one alternate allele, followed by the removal of background variation as described for the GATK VCF (ancestral sites with an MAF < 0.90 or sites in the ancestor that were either called as a variant or left uncalled). Next we applied hard filters to remove calls affected by mapping quality or strand bias (MQM > 40, SAF > 0 and SAR > 0, SAP > 0.5 and SRP > 0.5, RPR > 1 and RPL > 1). We then applied the same set of custom filters to the Freebayes VCF as described for the GATK VCF including the removal of sites with more than 10 missing samples, sites with an alternate allele count (AC) greater than three and sites with a Phred-based quality score (QUAL) below 200. Finally, based on the approximate read depth (DP) in this remaining set of variants we removed sites with more than 1.5 times the average to reduce false positives. To further reduce the number of false positives and misclassifications we manually reviewed this final set of SNPs using the Integrative Genomics Viewer (IGV) (Robinson *et al*. 2011, 2017).

### The Intersection of GATK and Freebayes VCFs and Variant Read Support

We viewed the two separately generated and filtered VCF files (from GATK and Freebayes) side-by-side and removed four sites that were not present in both files. The SNPs we removed included one SNP that had only been initially called by one of the callers and three SNPs that although initially called by both callers, only survived the filtration in one of the separate filtering pipelines. Going forward we worked with these cross-validated SNPs in the annotated GATK generated VCF.

As one final verification of variant support and quality we confirmed that each variant that had been called in a non-fruiting clone could also be detected in its origin population (or the line from which the clone was isolated). To do this, we generated read counts for each SNP across all samples using bam-readcount v0.7.4 (https://github.com/genome/bam-readcount). We removed any SNP for which the origin population did not support the same alternate allele with at least five reads, which we expect would only occur for false positives or for SNPs that did not provide a selective advantage. Also, because we expect that most true variants will be unique to one replicate line, for each SNP we viewed the number of reads supporting the same alternate allele across all samples. This led to the removal of one SNP that shared low level support for the same allele across multiple lines and also the rejection of one sample from a variant call (for details see *Variant Read Support* in the Supplemental Methods).

### Functional Annotation Clustering of Genes with SNPs

We performed functional annotation enrichment analysis with the final set of 29 genes containing SNPs using the online tool, DAVID (Database for Annotation, Visualization and Integrated Discovery (https://david.ncifcrf.gov/; Huang *et al*. 2009a, 2009b). We determined enriched terms using a Benjamini-corrected p-value (< 0.05) (Benjamini and Hochberg 1995).

### Structural Variant (SV) Calling and Filtration with Delly

We called structural variants (SVs) with Delly v0.8.3 (Rausch *et al*. 2012) following the Germline SV Calling workflow. We first did per-sample SV calling, providing the indexed, sorted and duplicate-marked BAM files and the indexed *D. discoideum* and *K. pneumoniae* concatenated reference genome. We then merged the SV sites to a single list and called genotypes across all samples. We merged all samples genotypes using bcftools v1.12 (using HTSLib v1.12) (Li 2011). We applied the Delly filter (-f germline) and then removed low quality variants that were not flagged as “PASS” in the VCF filter field (PE > 3 [or PE > 5 for translocations] and QUAL >=20). We annotated the SVs using the Ensembl Variant Effect Predictor (VEP 95.3) as described for the GATK VCF. We only retained simple, intrachromosomal SVs. As described for the SNPs, the same SV is unlikely to occur in more than one evolved line. But because we sequenced two or three samples for each line (the population and individual clones) the maximum number of times that a SNP is likely to appear in our data is twice (or three times for lines 20, 21 and 23). For this reason, we discarded SVs that were called in more than three samples. Last, as described for the SNPs, we manually reviewed this final set of SVs in IGV (Robinson *et al*. 2011).

### Association of Variants with the Loss of Altruism

To investigate for a potential correlation between variants and the complete loss of cooperation we returned to some of the evolved lines to isolate additional clones for targeted re-sequencing (Figure 1C). Each evolved line is a population comprised of both cooperators and evolved cheaters. The complete loss of cooperation is therefore only revealed when spores are plated clonally and can be identified by an inability to form a fruiting body (non-fruiting clones). Taking advantage of this population variation (Figure 2) we scored multiple additional clones for 1) whether or not it was able to form a fruiting body in isolation and 2) the presence or absence of the called variant(s) in the line.

For this analysis we again plated the evolved cell lines clonally (as described in the section, *Experimental Cell Lines*) to allow for growth from individual spores. After three to five days of growth we screened clones via PCR and Sanger sequencing. As previously described (in the section, *Experimental Cell Lines*), we took photographs of the screened clones to document the presence or absence of fruiting body formation (example images are available in Supplemental Material 1).

We generated genomic DNA for PCR genotyping clones using one of two methods. For some of the clones we first collected cells from the leading edge of the plaque, grew them to larger numbers and carried out a formal DNA extraction with the DNeasy® Blood and Tissue kit (Qiagen). To increase throughput, for most clones we directly lysed cells from the leading edge of the plaque and used the lysate for what we called a “plaque PCR.” Based on a protocol described by Charette and Cosson (2004), the plaque PCR included two steps. First, we used a sterile pipette tip to collect a small number of cells from the leading edge of a plaque and placed it in a tube containing 20 μl lysis buffer (10 mM Tris, pH 8.3, 50 mM KCl, 2.5 mM MgCl_2_, 0.45% Nonidet® P-40 (NP40) and 0.45% Tween® 20) with PK (1 μL of 20 μg/μL of PK for every 25 μL of lysis buffer). Next, we incubated the cells in lysis buffer for 1 min at 95° to inactivate the PK after which we used the cell lysate for PCR or stored them at −20°.

We designed primers to PCR-amplify the region spanning each called variant of interest (Supplemental Table S1). For the PCR we used either 1 μl of cell lysate or approximately 10 ng of DNA for the formal DNA isolations and the following reaction components: MgCl_2_ (25 mM) 1 µl, dNTPs (10 mM each) 0.5 µl, 5 pM of each primer, 5X GoTaq® Flexi Buffer 5 µl, GoTaq® DNA Polymerase 0.2 µl and H_2_0 10.3 µl for a 20 µl reaction. We used the following PCR protocol (adjusting the annealing temperature as needed, according to primer pairing): 95°, 2:00; 95°, 0:15; 50°, 0:15; 60°, 3:00; repeat steps 2–4 34X, 60°, 5:00; 4° hold. We submitted PCR products and primers (the same primers used for amplification) to Genewiz (South Plainfield, NJ) for purification and Sanger sequencing. We trimmed the returned sequences with 4Peaks (https://nucleobytes.com/4peaks/) for alignment using Seaview (http://doua.prabi.fr/software/seaview).

We screened a varying number of clones according to the availability of the fruiting and non-fruiting clones from each line. For each line included in this screen we tallied the number of clones that did and did not carry the called mutation(s) and whether or not the clone was able to form a fruiting body when plated clonally. We tested for significance using Fisher’s Exact Test with a 95% confidence interval using GraphPad Prism (version 9.3 for MacOS).

### Predicted Protein Structure

To explore proteins of interest, we first downloaded the amino acid sequences from DictyBase (Fey *et al*. 2009) and submitted them to Protter (Omasits *et al*. 2014) for visualization. We accessed three-dimensional structural predictions generated by AlphaFold (https://alphafold.ebi.ac.uk/) (Jumper *et al*. 2021; Varadi *et al*. 2022).

### Data Availability Statement

Raw reads in CRAM format will be stored in the Sequence Read Archive (SRA) [link/citation] (accession numbers/DOIs). Variant calls that survived filtration from each of the three variant callers (GATK, Freebayes and Delly) are available in Supplemental Material 2 in VCF format. All cell lines used in this study are in the lab of the senior authors.

## Results

### Raw Data Generation and Alignment

We obtained over 5 trillion reads across all samples, with a high rate of alignment (94.7%) to the concatenated *D. discoideum* and *K. pneumoniae* reference genomes. All reads that aligned to *K. pneumoniae* (food bacterium) were excluded from further analysis. The remaining 2.7 trillion reads aligned to *D. discoideum* with an average rate of 89.3%. Read alignment was, on average, equally successful for clones and evolved lines (89.3% and 89.4%, respectively). The average mapped read depth of all annotated genes normalized by gene length (gene annotations downloaded from NCBI on 10-25-19, Supplemental Material 3) is 54X and 195X for clones and evolved lines, respectively (Table 1). The average mapped read depth of normalized intergenic regions is very similar with averages of 53X and 180X for clones and evolved lines, respectively.

**Table 1.** Number of SNPs and average read depth by sample. Whole-genome sequencing and variant analysis resulted in 38 SNPs called in 25 of the 47 samples and distributed throughout 17 of the 24 evolved lines. This table indicates the number of SNPs called for each sample. The average mapped read depth was calculated for the raw reads normalized by the length of all annotated genes. Samples from each evolved line, sequenced as populations, are simply numbered 1 - 24. Clonal samples isolated from each of the evolved lines are named according to the population number hyphenated with the non-fruiting clone ID (e.g., “NF1”). The non-fruiting clone IDs were retained for record keeping purposes; they are not related to the number of clones sequenced for a line.

| Sample | SNP Count | Average Mapped Read Depth |
| --- | --- | --- |
| 1 | 0 | 128 |
| 1-NF2 | 0 | 43 |
| 2 | 1 | 137 |
| 3 | 0 | 146 |
| 3-NF1 | 1 | 127 |
| 4 | 0 | 171 |
| 5 | 0 | 161 |
| 5-NF2 | 3 | 40 |
| 6 | 1 | 152 |
| 7 | 4 | 229 |
| 7-NF2 | 5 | 44 |
| 8 | 0 | 165 |
| 8-NF1 | 0 | 89 |
| 9 | 1 | 220 |
| 9-NF3 | 1 | 24 |
| 10 | 1 | 151 |
| 10-NF3 | 1 | 23 |
| 11 | 2 | 116 |
| 11-NF1 | 0 | 37 |
| 12 | 1 | 215 |
| 12-NF1 | 1 | 97 |
| 13 | 1 | 165 |
| 13-NF2 | 1 | 70 |
| 14 | 0 | 127 |
| 14-NF3 | 0 | 22 |
| 15 | 0 | 176 |
| 15-NF1 | 0 | 69 |
| 16 | 0 | 277 |
| 16-NF2 | 3 | 118 |
| 17 | 3 | 123 |
| 18 | 0 | 191 |
| 18-NF1 | 3 | 30 |
| 19 | 0 | 319 |
| 19-NF1 | 0 | 80 |
| 20 | 0 | 231 |
| 20-NF1 | 1 | 66 |
| 20-NF3 | 1 | 32 |
| 21 | 1 | 272 |
| 21-NF1 | 3 | 40 |
| 21-NF2 | 5 | 45 |
| 22 | 3 | 291 |
| 23 | 0 | 305 |
| 23-NF1 | 0 | 25 |
| 23-NF2 | 0 | 43 |
| 24 | 0 | 222 |
| 24-NF1 | 1 | 32 |
| 1 | 0 | 128 |

### Resulting SNPs are Selection Driven

We called SNPs across all samples (joint calling) resulting in 53,220 unfiltered variants with GATK and 32,775 with Freebayes (before normalization). After independently filtering each call set the GATK VCF contained 306 SNPs and the Freebayes VCF contained 230 SNPs (the number of variants removed by each filter is available in Supplemental Material 4). We next manually reviewed all remaining SNPs in IGV resulting in approximately 75 sites in each call set. Finally, we removed any sites that did not co-occur in both call sets and/or sites lacking read support (as described in ***The Intersection of GATK and Freebayes VCFs and Variant Read Support***). The final set of SNPs contained 38 biallelic SNPs associated with 29 different genes (Table 2). Each gene in this list has only one SNP except for the gene, *grlG* (DDB_G0272244), which has 10 unique SNPs (discussed in detail in the following sections). All SNPs are unique to one evolved line with the exception of one in the unannotated gene, DDB_G0276529, which was called in two lines (7 and 16). Most evolved lines have between one and three SNPs except for lines 7 and 21, which have five and eight SNPs each, respectively (Table 1).

**Table 2.** List of called SNPs and the impacted locus. Whole-genome sequencing and variant analysis resulted in this list of 38 bi-allelic SNPs impacting 29 different genes. For each SNP this table provides the precise location in the genome and the Dictybase gene ID (followed by gene name when available), followed by the called samples. Samples from each evolved line, sequenced as populations, are simply numbered 1–24. Clonal samples isolated from each of the evolved lines are named according to the population number hyphenated with the non-fruiting clone ID (e.g., “NF1”). The last two columns of the table are the variant consequence and the estimated impact rating made by the Ensembl Variant Effect Predictor (VEP) (VEP 95.3).

| Chrom:Site | Gene ID | Called<br>Sample(s) | VEP Consequence | VEP<br>Impact |
| --- | --- | --- | --- | --- |
| NC_007087.3:2556092 | DDB_G0269332 | 22 | missense_var. | Moderate |
| NC_007087.3:3010110 | DDB_G0270828 | 21-NF1 | missense_var. | Moderate |
| NC_007087.3:3624901 | DDB_G0270964 | 18-NF1 | missense_var. | Moderate |
| NC_007087.3:3928085 | DDB_G0269956(RTE) | 21-NF2 | upstream_gene_var. | Modifier |
| NC_007087.3:4563335 | DDB_G0270834(*sgmA*) | 5-NF2 | missense_var. | Moderate |
| NC_007088.5:1741979 | DDB_G0272244(*grlG*) | 6 | stop_gained | High |
| NC_007088.5:1741982 | DDB_G0272244(*grlG*) | 7 & 7-NF2 | missense_var. | Moderate |
| NC_007088.5:1742043 | DDB_G0272244(*grlG*) | 11 | missense_var. | Moderate |
| NC_007088.5:1742355 | DDB_G0272244(*grlG*) | 17 | missense_var. | Moderate |
| NC_007088.5:1742362 | DDB_G0272244(*grlG*) | 13 & 13-NF2 | missense_var. | Moderate |
| NC_007088.5:1742375 | DDB_G0272244(*grlG*) | 10 & 10-NF3 | stop_gained | High |
| NC_007088.5:1742506 | DDB_G0272244(*grlG*) | 18-NF1 | missense_var. | Moderate |
| NC_007088.5:1743478 | DDB_G0272244(*grlG*) | 2 | missense_var. | Moderate |
| NC_007088.5:1743823 | DDB_G0272244(*grlG*) | 21 & 21-NF2 | stop_gained | High |
| NC_007088.5:1743844 | DDB_G0272244(*grlG*) | 22 | stop_gained | High |
| NC_007088.5:1761939 | DDB_G0272484 | 16-NF2 | missense_var. | Moderate |
| NC_007088.5:4313995 | DDB_G0274875(*rnf160*) | 16-NF2 | stop_gained | High |
| NC_007088.5:6565220 | DDB_G0276291 | 17 | missense_var. | Moderate |
| NC_007088.5:6762467 | DDB_G0276367 | 17 | stop_gained | High |
| NC_007088.5:6787007 | DDB_G0276529 | 7-NF2 & 16-NF2 | missense_var. | Moderate |
| NC_007088.5:6871012 | DDB_G0276553 | 7 & 7-NF2 | upstream_gene_var. | Modifier |
| NC_007088.5:7954466 | DDB_G0277481 | 21-NF2 | missense_var. | Moderate |
| NC_007089.4:1029309 | DDB_G0278531 | 24-NF1 | stop_gained | High |
| NC_007089.4:1073554 | DDB_G0278559 | 7 & 7-NF2 | missense_var. | Moderate |
| NC_007089.4:1101637 | DDB_G0278575 | 7 & 7-NF2 | upstream_gene_var. | Modifier |
| NC_007089.4:3459101 | DDB_G0280505(*tmem144A*) | 5-NF2 | missense_var. | Moderate |
| NC_007089.4:5126190 | DDB_G0281923(*mrhA*) | 21-NF1 | downstream_gene_var. | Modifier |
| NC_007089.4:5671190 | DDB_G0282355 | 12 & 12-NF1 | upstream_gene_var. | Modifier |
| NC_007090.3:2549009 | DDB_G0284845(*gxcC*) | 3-NF1 | upstream_gene_var. | Modifier |
| NC_007091.3:1993104 | DDB_G0288805 | 22 | stop_gained | High |
| NC_007091.3:2942727 | DDB_G0289555(*arkA*) | 21-NF1 | missense_var. | Moderate |
| NC_007091.3:2996049 | DDB_G0289583 | 11 | missense_var. | Moderate |
| NC_007091.3:4205385 | DDB_G0290523 | 21-NF2 | synonymous_var. | Low |
| NC_007091.3:4793752 | DDB_G0290943(*pks39*) | 20-NF1 & 20-NF3 | missense_var. | Moderate |
| NC_007091.3:4956884 | DDB_G0291085(*gxcE*) | 21-NF2 | missense_var. | Moderate |
| NC_007091.3:905736 | DDB_G0287967 | 18-NF1 | downstream_gene_var. | Modifier |
| NC_007092.3:1299121 | DDB_G0292198(RTE) | 5-NF2 | missense_var. | Moderate |
| NC_007092.3:1984519 | DDB_G0292696(*colA*) | 9 & 9-NF3 | missense_var. | Moderate |

Most SNPs (31 of 38) are in coding regions of the genome and all but one of these result in an introduced stop codon or amino acid substitution (Table 2). Among the 31 SNPs in coding sequence, 22 are missense variants and according to the Variant Effect Predictor annotations are predicted to have a moderate impact, eight others introduce a premature stop codon with a high predicted impact and the one synonymous variant has a low predicted impact. Among the seven SNPs in non-coding sequence, five are upstream and two are downstream variants that the Variant Effect Predictor annotated as “modifier,” meaning they are either difficult to predict or that there is no evidence of an impact. This distribution of large effect SNPs strongly supports that they have increased in abundance due to selection, rather than drift.

### Genes with SNPs

The 29 genes with SNPs (Table 2) are distributed throughout the *D. discoideum* genome with between one and eight SNPs on each of the six chromosomes. The average GC content for the set of 29 genes (excluding introns) is 28.4% which is close to the genome-wide average of 27% for protein-coding genes. Each of the genes in our list has one SNP except for the G protein-coupled receptor (GPCR), *grlG*, which has 10 unique SNPs. *GrlG* is one of 17 glutamate receptor-like (’grl’) proteins in the *D. discoideum* genome (*grlA*–H and *grlJ*–R) (Eichinger *et al*. 2005) but the function of *grlG* is not yet known. And like the majority of glutamate receptor-like proteins in *D. discoideum*, we do not know the ligand(s) that it binds nor the downstream effector(s). Among the remaining 28 genes (each with one unique SNP) a few have been well described (*e.g., sgmA*, *pks39* and *arkA*) but the majority (17) are hypothetical proteins still largely lacking annotation.

Using annotations that are available for our 29 genes with SNPs, we carried out functional annotation clustering with DAVID and identified two significantly enriched annotation clusters. The first cluster of five annotations contained UniProt Keywords (Bateman *et al*. 2021) related to zinc and metal binding (group enrichment score of 1.45) including *arkA*, DDB_G0272484, *rnf160*, DDB_G0269332 and *sgmA*. The second cluster of 10 annotations contained terms related to transmembrane and membrane annotations (group enrichment score of 1.11). This second cluster contained *arkA* and DDB_G0269332 from the first cluster as well as *grlG*, *pks39*, DDB_G0276291, DDB_G0278531, DDB_G0278575, DDB_G0281923, DDB_G0290523 and *tmem144A*. After applying the Benjamini (Benjamini and Hochberg 1995) correction, none of the individual terms are significantly enriched (Supplemental Material 5). The large number of unannotated genes implicated here does hinder interpretation, but it is not unusual given that at the time of writing this manuscript, approximately 40% of protein-coding genes in the *D. discoideum* genome still lack annotation (Fey *et al*. 2009).

### Structural Variants (SVs) Provide Further Support for grlG

We called structural variants (SVs) across all samples (joint calling) using Delly resulting in an unfiltered set of 10,139 SVs. The Delly filter reduced the number to 131 and after all quality filtration we have a set of 12 SVs (Supplemental Table S2). Among the 12 SVs are six deletions, five inversions and one duplication. Nine of the 24 evolved lines carry 1 or more of these SVs, two of which (lines 1 and 14) did not have any called SNPs. Each SV is unique to one evolved line with the exception of one deletion (NC_007088.5:1742215–1742506) in *grlG* which was called in lines 17 and 18. However, because we cannot rule out potential contamination of the population sample of line 18 we only report the variant in line 17 (more detail is available in *Variant Read Support* in the Supplemental Methods). The deletion was verified by PCR in line 17 (as part of the analysis described in the section, *Association of Variants with the Loss of Altruism*). The SVs range considerably in length (from 183 bp to 49 Kb) potentially impacting as many as 21 different genes. Strikingly, five of the 12 SVs impact *grlG*, the same gene that we already identified as carrying 10 SNPs. None of the 28 other genes with SNPs were impacted by any of the SVs. Given the clear signals of parallel evolution identified for *grlG*, we chose not to pursue the other SVs any further. We will only discuss the SVs that impact *grlG* going forward; the full set of 29 SVs that passed filtration are available in VCF format in Supplemental Material 2.

### Some Variants in grlG are Associated with the Loss of Cooperation

Given the high level of parallelism identified in *grlG* (one or more variants in more than half of the evolved lines) we returned to the evolved lines to screen additional fruiting and non-fruiting clones for a correlation between variants in *grlG* and the loss of cooperation. The variants are located throughout the length of the gene but a pattern emerged in which only those variants located in the 5’ half of *grlG* (on exon 1) are associated with the loss of cooperation (*i.e.,* non-fruiting clones) and variants in the 3’ half (on exon 2) are not (Figure 3 and Supplemental Table S3). Because *grlG* has two exons of roughly equal length (383 and 390 amino acids for exon 1 and exon2, respectively) and for simplicity, we will refer to the two halves as the 5’ and 3’ regions.

**Figure 3.**
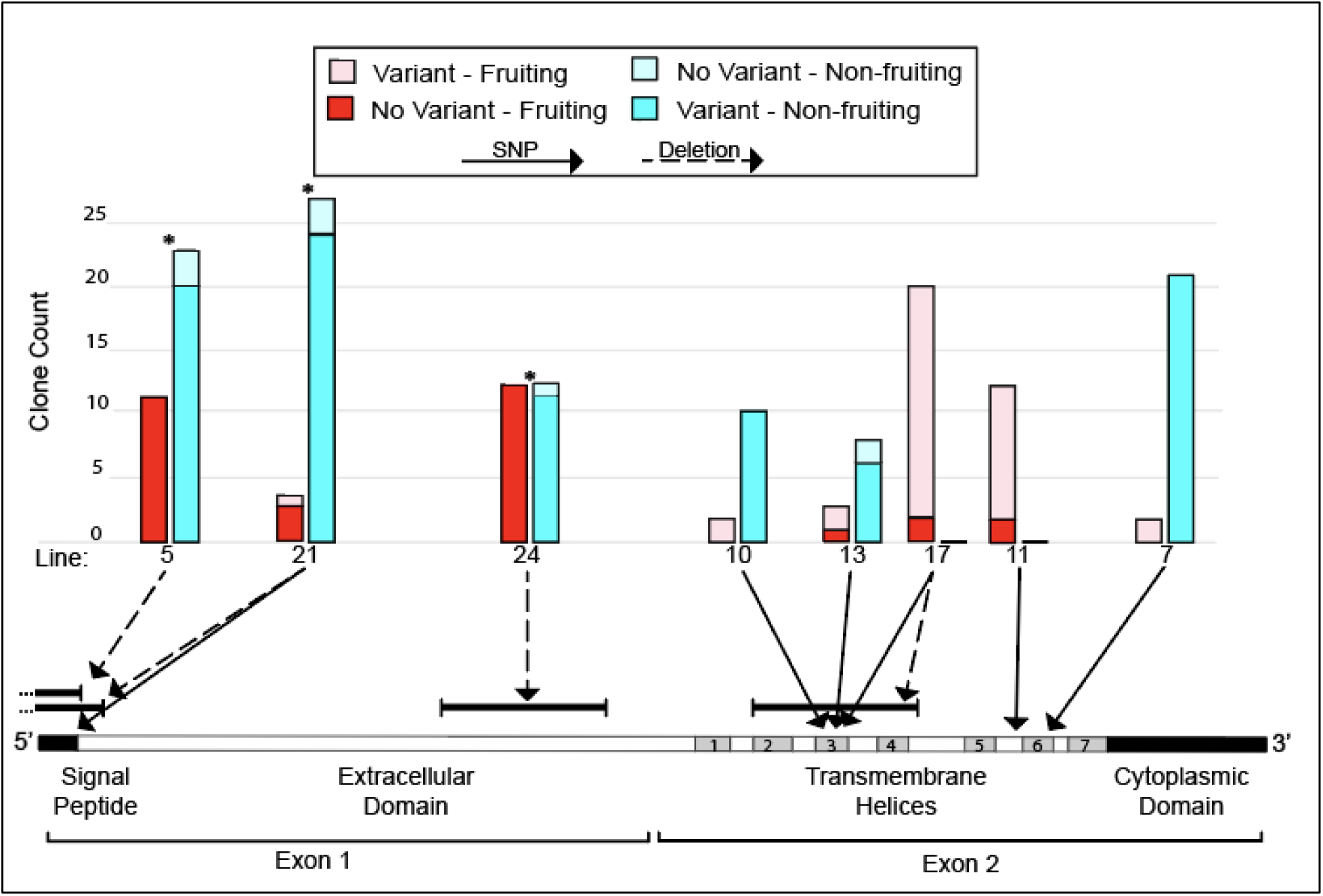
Location of variants in *grlG* and correlation with the loss of cooperation. Schematic representation (drawn to scale) of the coding sequence of *grlG*. The major features of a mature *grlG* protein are indicated below the schematic (see also, Figure 5). Ten of the 15 called variants (labeled above the schematic) were included in a targeted resequencing of additional clones from each respective evolved line to investigate the association between variants in *grlG* and the loss of cooperation (clonal inability to form a fruiting body). For each evolved line that we screened clones from, a histogram illustrates the results with side-by-side bars showing the number of clones that could (left bar) and could not (right bar) form a fruiting body in isolation and the number of each phenotype (fruiting or non-fruiting) that did or did not carry the called variant. Significant associations are indicated by an asterisk (*).

We prioritized screening of variants that we had identified in evolved lines with a moderate percentage of non-fruiting clones (Figure 2) so that both fruiting and non-fruiting clones could be included. We also prioritized variants with moderate allele frequencies in their evolved populations. As such, the analysis included ten of the 15 variants identified across the length of *grlG*. The 10 variants were called in 8 different lines (two unique variants were called in lines 21 and 17), three of which have variants in the 5’ region (lines 5, 21 and 24) and five with variants in the 3’ region (7, 10, 11, 13 and 17). From the three lines with variants in the 5’ region, we screened 88 clones (62 non-fruiters and 27 fruiters) for the presence of four called variants. We found that 88.7% (55 of 62) of the non-fruiter clones carried the variant compared to only 3.7% (1 of 27) of the fruiting clones. The association between the presence of variants in the 5’ region of *grlG* and the clonal phenotype is significant in each evolved line we screened (Fisher’s Exact p < 0.0001, p = 0.0164 and p < 0.0001 for lines 5, 21 and 24, respectively). In the 3’ region, from the five lines with variants, we screened 78 clones (39 non-fruiters and 39 fruiters) for the presence of six called variants. Unlike the association in the 5’ region, we found the number of clones carrying a variant in the 3’ region was roughly equal between non-fruiters and fruiters (94.8% [37 of 39] and 87.2% [34 of 39], respectively). There is no association between the presence of variants in the 3’ region of *grlG* and the clonal phenotype in any of the evolved lines we screened (Fisher’s Exact p > 0.05).

According to our results, variants in the 5’ region of *grlG* are significantly associated with the non-fruiting phenotype but not all clones follow that trend (Figure 3). In all three evolved lines that we screened in the 5’ region, we found a few clones that did not have a variant, but they were still unable to form fruiting bodies in isolation. These clones could have lost cooperation because of a different, undetected variant (anywhere in the genome). However, a clone that is still able to cooperate and form a fruiting body in isolation, despite the apparent loss of *grlG,* is more difficult to align with our hypothesis that high impact variants in *grlG* cause non-fruiting. We only found one such clone in the 5’ region. One clone from line 21 was able to form a fruiting body despite carrying a SNP that introduces a premature stop codon in *grlG* after only 23 amino acids. None of the other 26 fruiting clones that we screened in the 5’ region carried a variant in *grlG*. This is strikingly different from the results in the 3’ region, where almost all of the fruiting clones that we screened carried a variant in *grlG,* though these were concentrated in lines 11 and 17 and rare in others. Thus, variants in the 5’ region of *grlG* usually result in the loss of cooperation but variants in the 3’ region do not. For a subset of the clones that we screened for *grlG* variants we also screened for the presence of one or more SNPs that we called in other gene(s). These additional screens were largely uninformative and are therefore only described in the supplemental materials (Additional File 1 and Supplemental Table S4).

### The Genes Discovered in This Study Have Not Previously Been Implicated in Cheating

To look for a relationship between the 29 genes identified herein and the loss of altruism, we turned to the literature to see if any of them have been previously implicated in cheating in *D. discoideum*. Because we are looking for obligate cheater genes, most notably missing from our list is *fbxA,* the only previously known obligate cheater gene (Ennis *et al*. 2000). It is also interesting that our list does not contain any of the characterized facultative cheater genes such as *chtB, chtC*, *dimA* or *csA* (Queller *et al*. 2003; Foster *et al*. 2004; Khare and Shaulsky 2010; Santorelli *et al*. 2013). Aside from that handful of genes that first come to mind, the remaining majority of genes with potential involvement in cheating in *D. discoideum*, have not yet received attention at the individual level. We were particularly interested in the many loci recovered in the large restriction enzyme-mediated insertional (REMI) mutant library screening from Santorelli *et al*. (2008). In that study, a set of 167 mutants (corresponding to 197 loci) were identified that increased under low-relatedness selection. Based on testing a subset of those mutants, the authors estimate that approximately 75% of the mutants are cheaters, but so far, only *chtB* and *chtC* have been described. Although Santorelli *et al*. (2008) focused on facultative cheater genes (by requiring one round of solo fruiting body formation) they did recover *fbxA* and a few other mutants were found with morphological abnormalities indicating there may be a others that are not facultative. The full set of 197 loci identified by Santorelli *et al*. (2008) is the first of four gene lists we chose for cross-comparison. The other gene sets are each comprised of genes found to be differentially expressed in chimeric mixtures identified at various stages of multicellular development and thus may contain genes involved in conflict and cheating. Two are comprised of genes identified at tight aggregate stage (79 genes) or the slug stage (190 genes) (Noh *et al*. 2018 and Oliveira *et al*. 2019, respectively). The last gene set is composed of 152 genes that were also identified during the establishment of multicellularity (at either 8 or 12 hours), but in this case, the strains had been engineered to be mutually incompatible (*tgrB1*/*tgrC1* gene replacement strains) which may increase the potential to identify genes involved in conflict (Hirose *et al*. 2015). Unlike the genes from Santorelli et al. (2008) these last three gene sets have not been directly implicated in cheating. But given their differential expression in chimeras, a time at which cheating or the resistance to cheating would be adaptive (Noh *et al*. 2018), they are likely to include genes involved in conflict and cheating. In support of this reasoning, the 79 genes from Noh *et al*. (2018) show signatures of rapid molecular evolution, consistent with conflict driven escalating arms races (Queller and Strassmann 2018; Scott and Queller 2019). For simplicity, we will refer to these four sets of genes as candidate cheater genes.

Altogether, the four sets of candidate cheater genes from the literature comprise a list of 591 unique genes (Supplemental Table S4). None of the 29 genes identified in this study were found on that list and there is little overlap among the four candidate cheater gene sets (Figure 4). This in line with previous reports of limited overlap among sets of social genes (kin discrimination, cooperation and cheating) (Oliveira *et al*. 2019; Noh *et al*. 2020). Only 26 of the 591 genes are present on more than one list. No genes are present on all four lists and only one gene is present on three of the four; the pre-spore specific gene D (*pspD*). Because the genes from Santorelli *et al*. (2008) were identified as part of a screen for cheater genes specifically, they are of particular interest. Eleven genes from Santorelli *et al*. (2008) were found on one or more of the other gene lists and three of them were among the subset of mutants directly tested for cheating in that study. One of these tested mutants (LAS92) had an insertion in a gene that is also on the list from Hirose *et al*. (2015) and it was confirmed to be a cheater (DDB_G0283373). Two other tested mutants had intergenic insertions near genes that are also on the list from Oliveira *et al*. (2019); one of which was confirmed to be a cheater (LAS6) (DDB_G0276335) and the other was not (LAS60) (DDB_G0285843). However, these last two mutants from Santorelli *et al*. (2008) had intergenic insertions and the actual gene(s) impacted was not precisely determined.

**Figure 4.**
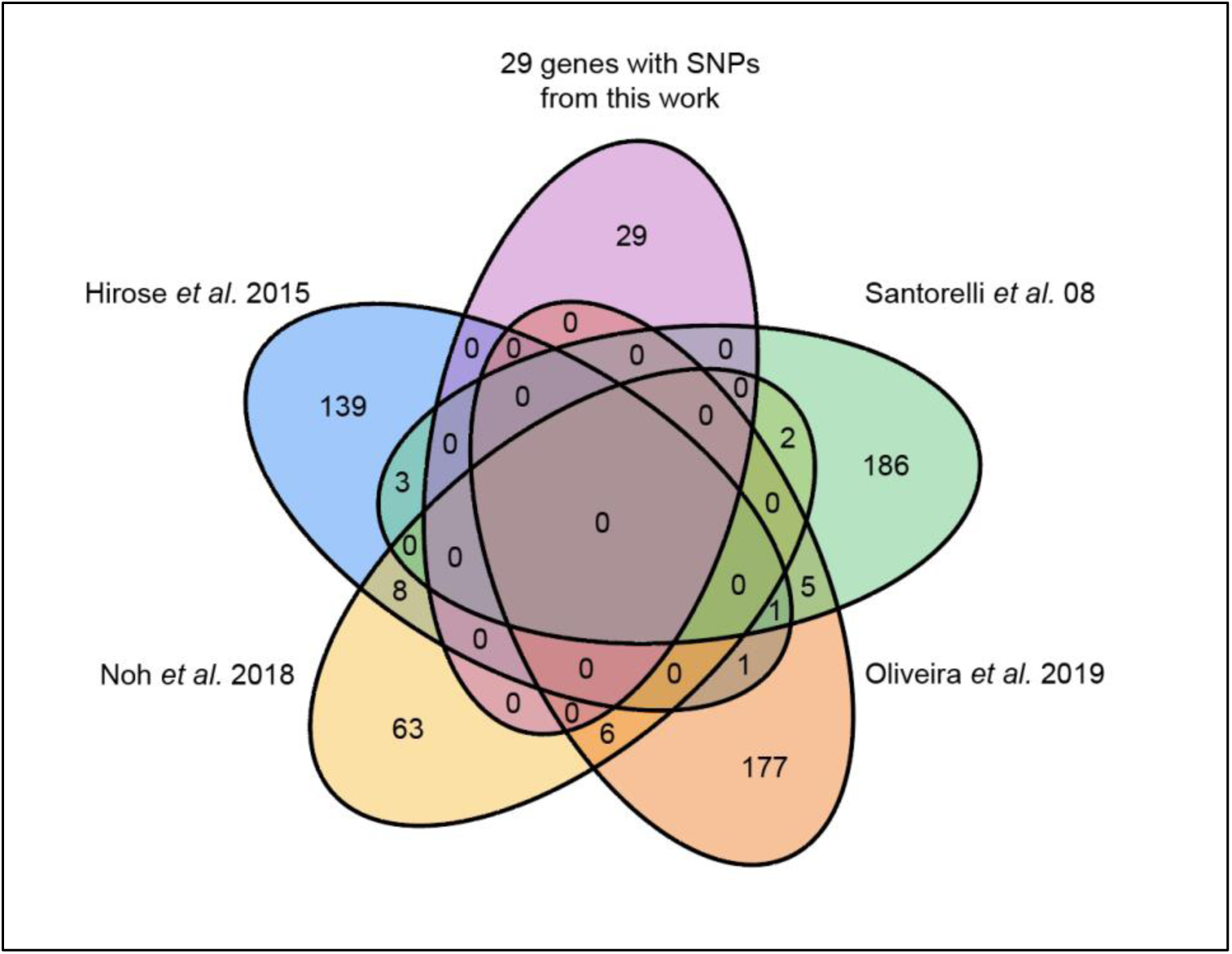
Limited overlap among published candidate cheater gene sets. We compared our list of 29 genes with SNPs to four genes sets from the literature including cheater genes and genes differentially expressed in chimera during the social stage (potentially involved in cheating). Together, the full list is comprised of 591 genes that have been implicated in cheating or cooperation (Santorelli *et al*. 2008; Hirose *et al*. 2015; Noh *et al*. 2018; Oliveira *et al*. 2019). Each gene set is shown in a different color and outlined with a dashed line. Overlapping portions represent the number of genes shared among the different gene sets. The total number of genes for each set is as follows: 197 from Santorelli *et al*. (2008), 152 from Hirose *et al*. (2015), 79 from Noh *et al*. (2018) and 190 from Oliveira *et al*. (2019). None of the 29 genes with SNPs identified in this work were found among the gene sets from the literature and there is very little overlap among the other four lists.

## Discussion

In this study we identified SNPs in 29 genes that arose across 24 replicate lines of *D. discoideum* experimentally evolved by Kuzdzal-Fick *et al*. (2011) under conditions that mean low-relatedness for any new mutations. In that earlier study, over the course of about 290 generations at low-relatedness, 19 of the 24 replicate lines saw a significant rise in cheating, defined as a mutant contributing proportionally more to spore cells than stalk cells. When isolates from these lines were plated clonally, an average of 31% had completely lost altruism and were unable to form fruiting bodies on their own. Because each line is a mixed population, the mutations that conferred the obligate cheating advantage were able to spread by forming chimeric fruiting bodies with cooperators lacking the mutation. Here we used whole-genome sequencing and variant analyses of those previously evolved cell lines to identify which genes had changed and might be responsible for the non-fruiting, obligate cheating phenotype. While there are many genes known to cause facultative cheating, before this study only one gene (*fbxA*) had been recognized to cause obligate cheating. We identified at least one SNP in 17 of the 24 experimentally evolved lines for a total of 38 SNPs. Most of the SNPs (31) are in coding regions of the genome, and all but one results in either an introduced stop codon or other amino acid substitution suggesting selective pressure acting on these loci under the experimental conditions in Kuzdzal-Fick *et al*. (2011). The majority of the genes with SNPs are not well characterized and many lack any annotation which impacted our ability to assess any uniting similarities among them. Except for one of these genes, *grlG*, with 10 unique SNPs across multiple lines, the other 28 genes only have one SNP each. *GrlG* also stands out because we found it also harbors five structural variants, whereas none of the other genes with SNPs had any. Our results provide strong evidence of parallel evolution of *grlG* and suggests the involvement of this G protein-coupled receptor (GPCR) in cooperation and multicellular development in *D. discoideum*.

### Confidence in Variant Calls and Detection

The *D. discoideum* genome has features that may have impacted our ability to detect variants. The AT content of the *D. discoideum* genome is very high, greater than 77% (Eichinger *et al*. 2005). The high AT content contributes to the high density of simple sequence repeats (DNA tracts of 1–6 bp tandemly repeated a varying number of times) (Tian *et al*. 2011) which represent 14.3% of the genome (Srivastava *et al*. 2019). Moreover, in *D. discoideum*, these repeats are not confined to non-coding regions; they are also found in over 16.3% of protein coding genes (Eichinger *et al*. 2005). Simple sequence repeats are recognized as mutational hotspots across a number of organisms and are particularly prone to insertions and deletions (indels) (Tóth *et al*. 2000; Ellegren 2004). In *D. discoideum*, both indels and SNPs have been shown to occur at a higher frequency in low-complexity, repetitive regions of the genome (Srivastava *et al*. 2019). The precision of variant calling is also decreased in the same sorts of low-complexity and repeat regions, compounding the challenges of variant calling in *D. discoideum*. This decreased precision is particularly true for detection of structural variants, which are inherently more challenging to identify, particularly with short-read sequence data (Cameron *et al*. 2019; Kosugi *et al*. 2019; Mahmoud *et al*. 2019). Despite these challenges, we chose not to exclude low-complexity or non-coding regions of the genome, which could lead to undiscovered variation outright. Instead, we dealt with the challenging features of the *D. discoideum* genome by carefully exploring the effects of different settings for variant calling and filtering to determine a stringent and robust pipeline. We separately called SNPs with two independent callers. We applied our carefully chosen quality filters to the two call sets independently, followed by manual inspection of all variants. We then conservatively only pursued the SNPs still present in both call sets. Finally, we used the BAM read counts across all samples to rule out potential contamination or other false positives. Together, despite the challenging features of the *D. discoideum* genome, we are confident that we obtained a high quality and strongly supported set of variants.

Our stringent pipeline decreased the likelihood of obtaining false positives and ensured that variants called at the population level tend to be those most strongly selected. The trade-off of such a stringent approach is a decreased sensitivity to detect low frequency variants in the population samples. Given the relatively short amount of time the experimental lines were allowed to evolve (Kuzdzal-Fick *et al*. 2011), not all variants would have had the time to expand throughout the population of that line. Therefore, variants that arose later in the experiment or those not facing strong selection, may have been difficult for us to detect at the population level. Although we cannot rule out the possibility that some were left undetected, we did detect some low frequency variants by sequencing individual non-fruiting clones from the lines.

### Parallel Evolution and the Involvement of grlG in Cooperation

The concentration of highly supported and unique variants in *grlG* across 14 of the 24 replicate lines is strong evidence of parallel evolution at the gene level (Tenaillon *et al*. 2012; Barrick and Lenski 2013; Van den Bergh *et al*. 2018). None of these mutations were present in the ancestral line, indicating that they occurred during the course of experimental evolution (Kuzdzal-Fick *et al*. 2011). At the nucleotide level there is no indication of a mutational hotspot because the variants are located throughout the length of the gene. Moreover, the variants in *grlG* are comprised of several types of mutations including 10 non-synonymous SNPs that result in either an introduced stop codon (four) or a missense mutation (six), as well as four deletions and one inversion, all of which impact coding sequence. The number of large-effect mutations (half of which result in protein truncation) and the complete lack of synonymous mutations in *grlG* suggests that the loss of *grlG* was adaptive under the experimental conditions. Further support of selection acting on *grlG* is the high population level frequency that several of these new mutations reached in their respective lines. On average, the 10 unique SNPs in *grlG* are supported by 61% of the total reads in their respective populations compared to an average of only 41% support for the 28 remaining SNPs in other genes. These data suggest that *grlG* variants increased in abundance due to selection, rather than drift. In addition, by screening multiple additional clones from the experimental lines, we found that variants in the 5’ region of *grlG* are significantly associated with the loss of cooperation (non-fruiting) while those in the 3’ region are not. However, very few of the clones screened in the 3’ region carried a variant in *grlG* (9%) and this may have limited our power to detect an association. We were unable to find any non-fruiting clones for evolved lines 11 and 17 but the large proportion of normally fruiting clones that carry a *grlG* variant certainly suggests a lack of an association in those lines. To help us make sense of these results, we next explore the limited amount of currently available information about *grlG* and view our findings in light of what is known more broadly about the large and highly conserved family of proteins, the G protein-coupled receptors (GPCRs).

The G protein-coupled receptors (GPCRs) in eukaryotes are the largest class of receptors of extracellular stimuli. They generally activate intracellular responses via heterotrimeric G-proteins. In humans there are over 800 GPCR genes making them the most abundant gene family in the human genome (Lagerström and Schiöth 2008). They are involved in a huge diversity of biological processes including signaling of hormones, metabolites and neurotransmitters to cell migration and development (Rosenbaum *et al*. 2009). The GPCRs are responsible for approximately 66 known single-gene human diseases, and they are the target of about one third of FDA-approved drugs (Hauser *et al*. 2017; Schöneberg and Liebscher 2021). Thus, identifying the ligands and the roles of yet uncharacterized GPCRs (or, orphan GPCRs) is of great interest broadly and the GPCRs in *Dictyostelium* are no exception. *Dictyostelium discoideum* is an established model system for studying numerous processes such as chemotaxis and other cell-motility linked processes such as phagocytosis and development (Bozzaro 2019), all of which involve GPCRs.

The *D. discoideum* genome has a surprisingly large and diverse repertoire of more than 55 GPCRs (Eichinger *et al*. 2005; Sucgang *et al*. 2011; Heidel *et al*. 2011). Classically the GPCRs are categorized into six classes based on sequence and function (Kolakowski 1994; Bockaert and Pin 1999); *D. discoideum* has representative members of most classes (Prabhu and Eichinger 2006). Similar to those in vertebrates, the GPCRs in *D. discoideum* are involved in a diversity of biological processes, including significant roles in development. Most notably, members of the class E GPCRs include the receptors of cAMP (cARs). The chemoattractant cAMP is pivotal in the initiation and coordination of multicellular development in *D. discoideum* (Saran *et al*. 2002; Loomis 2014). Perhaps it is not surprising then that the cAMP receptors were the first GPCRs to be identified (Klein *et al*. 1988) in *D. discoideum* and remain the most extensively studied.

Some members of the class C GPCRs (which includes *grlG*) in *D. discoideum* have also been implicated in development. The *D. discoideum* genome has 17 members of the class C GPCRs, known as glutamate receptor-like (grl) proteins (Eichinger *et al*. 2005). For most of the group, the ligands, effectors and even the specific signaling pathways remain unknown. A great deal of progress has been made over the past few decades especially after the completion of the *D. discoideum* genome in 2005 (Eichinger *et al*. 2005). At least four of the glutamate receptor-like proteins have been described as having roles in development including *grlA* (Prabhu *et al*. 2007a; Anjard *et al*. 2009), *grlB* (Wu and Janetopoulos 2013), *grlE* (Anjard and Loomis 2006; Wu and Janetopoulos 2013) and *grlJ* (Prabhu *et al*. 2007b). Moreover, Prabhu *et al*. (2007b) also found that all 17 are expressed throughout development, some with increased expression early in aggregation (*grlA, D, E, J, M, N, Q*) and others with increased expression from the tight aggregate stage and on (including *grlG, B, C, F, H, K, L, O, P, R*). Much remains to be learned but it is easy to conceive that *grlG,* like other members of this protein family in *D. discoideum*, is also involved in cooperation during multicellular development.

We are aware of only one previous study that investigated the function of *grlG* but it was in the context of predation rather than development. Upon exposure to the chemoattractant folic acid (used by *D. discoideum* to chemotax toward and phagocytose bacteria) Pan *et al*. (2016) identified increased phosphorylation at key serine residues in both *grlG* and *grlL.* Both *grlG* and *grlL* were thus considered candidate folic acid receptors (they referred to them as folic acid receptors (’far’) *far2* and *far1,* respectively). However, only *far1* (*grlL*), and not *grlG,* was required for eliciting the response to folic acid and the authors moved forward with the characterization of only *far1* (*grlL*) (Pan *et al*. 2016, 2018). During the work that followed, they noticed that the loss of *far1* (*grlL*) does not completely abolish the response to folic acid and lipopolysaccharide (LPS) which suggests that other proteins are involved. That observation is in line with some earlier studies which also suggested that there are two folic acid receptors, each with a different binding affinity (de Wit and van Haastert 1985; Segall *et al*. 1988). But *grlG* is not required for the folic acid response and there is only limited evidence to support any involvement (Pan *et al*. 2016; Lamrabet *et al*. 2020). We have no reason to believe that the parallel evolution of *grlG* that we have identified is related to the detection or phagocytosis of bacterial prey. The *D. discoideum* AX4 lines were adapted to the laboratory growth and nutrient conditions well before the start of experimental evolution by Kuzdzal-Fick *et al*. (2011). Instead, the primary novelty of the experimental environment was the extreme low-relatedness that favored cheaters.

The number of large-effect variants in *grlG* suggests that the loss of *grlG* was adaptive under the experimental conditions of low relatedness. What is less clear from our results is why only the variants in the 5’ region of *grlG* are significantly associated with non-fruiting and those in the 3’ region are not. To investigate, we viewed the amino acid sequence topology and predicted protein structure to investigate other potential explanations (Figure 5B). *GrlG* shares the same major features found in most class C GPCRs including a long 5’ extracellular domain, a seven-transmembrane domain toward the 3’ end followed by an intracellular C-terminal tail (Figure 5B) (Pin *et al*. 2003; Rosenbaum *et al*. 2009). Ligand binding in class C GPCRs typically occurs via a Venus flytrap domain (similar to the ancestral periplasmic binding proteins of bacteria) situated in the 5’ extracellular domain (Cao *et al*. 2009; Chun *et al*. 2012). Upon binding its ligand, conformational changes throughout the seven-transmembrane domain lead to the activation of G proteins (or other effectors) at the intracellular C-terminus to induce signaling inside the cell (Rosenbaum *et al*. 2009). Based on sequence homology and the predicted folding structure, *grlG* appears to function similarly. In the predicted structure generated by AlphaFold (Jumper *et al*. 2021; Varadi *et al*. 2022), the extracellular region can be seen folded (with high confidence) into a Venus flytrap structure with a clearly visible cleft, or ligand binding pocket (Figure 5A).

**Figure 5.**
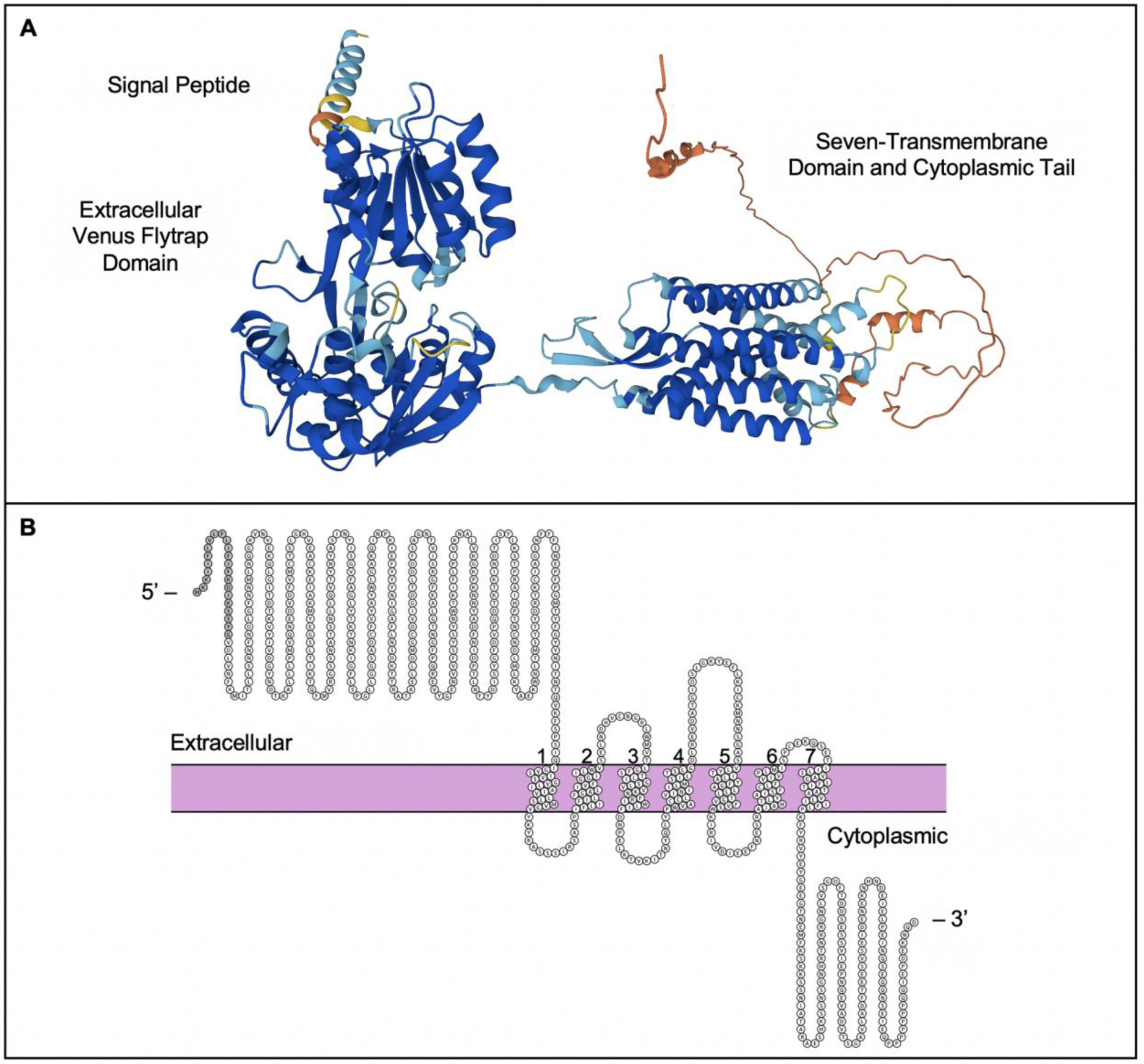
Predicted structure and sequence topology of *D. discoideum grlG.* **A**. Predicted folding structure of *grlG* generated by AlphaFold (Jumper *et al*. 2021; Varadi *et al*. 2022). Colors indicate the model confidence in the local accuracy; dark blue is very high confidence, light blue is confident, yellow is low confidence and orange is very low confidence. This structure can be be viewed interactively with the 3D viewer on the AlphaFold Protein Structure Database (https://alphafold.ebi.ac.uk/entry/Q75JP4). **B.** The sequence topology of *grlG* generated in Protter (Omasits *et al*. 2014). Extracellular regions are shown above the membrane (in purple) and the cytoplasmic regions below. The helices of the seven-transmembrane domain are numbered one through seven. The 5’ amino acids in grey indicate the signal peptide.

According to the amino acid sequence topology, each of the four variants that we screened for in the 5’ region would result in early protein truncation, either upstream of or within the extracellular binding domain. It is not surprising that the loss of the binding domain, such as in our 5’ variants, would render a receptor non-functional. Similar results have been experimentally demonstrated for the homologous protein, *grlL* (*far1*). Pan *et al*. (2018) generated a *grlL* (*far1*) mutant missing only the extracellular binding domain (the remainder of the protein was intact) to demonstrate that the mutant was unable to elicit a response to its ligands. But while we might predict that the large impact variants in the transmembrane domain, particularly those also resulting in early protein truncation, would lead to similar outcomes, our data do not support a significant association between any of the 3’ variants and non-fruiting. Additional work is needed to understand how *D. discoideum* cells handle aberrant *grlG* mRNAs and proteins. But even if we assume that aberrant and truncated proteins are not degraded, given the coordinated functionality between extracellular ligand binding and the resultant intracellular signaling, the loss of either end of this transmembrane protein receptor would seem to disable signaling. Moreover, the variants in both regions of *grlG* were positively selected and increased in abundance under the experimental conditions. Although it is difficult to speculate given so many unknowns, perhaps all variants in *grlG* (both 5’ and 3’) provided the selective advantage of cheating, but only those that inhibit ligand binding are also likely to result in non-fruiting when in isolation.

Another challenge to interpreting our results is that variants in *grlG* are apparently not completely penetrant. While our results are broadly consistent with variants in 5’ *grlG* causing non-fruiting, we did identify one clone in line 21 that was still able to form a fruiting body despite carrying a variant in 5’ *grlG* that introduces a premature stop codon just before the extracellular binding domain. Although human or technical error is a possibility, there also many potential biological explanations. Some of the factors that can influence penetrance include gene expression levels, modifier genes, the co-occurrence of one or more additional variants, the environment or other non-genetic factors (Cooper *et al*. 2013). However, the lack of penetrance we observed may not be explained until we know more about the gene, particularly the ligand(s) it binds and the pathway(s) it is involved in. This brings our attention to the complex and pleiotropic nature of GPCR signaling and highlights the challenge of predicting variant impacts.

From the great deal of work with GPCRs in humans and disease in particular, it is clear that mutations in membrane proteins can result in a huge variety of impacts. The effect of any mutation will depend on the properties of the protein as well as the type and location of the mutation (Schöneberg and Liebscher 2021; Zaucha *et al*. 2021). Some of the many potential impacts of GPCR mutations include improper membrane trafficking or localization; alterations to critical secondary or tertiary structures; inhibition or altered efficiency of binding ligand(s) or interacting proteins and inhibition or altered efficiency of signaling through downstream effectors. Further complicating matters, some GPCRs bind multiple ligands and/or have multiple effectors. Likewise, one ligand can have multiple receptors. In *D. discoideum* for example, the chemoattractant cAMP has four receptors (cARs 1–4) in the GPCR class E, each of which has a different cAMP-binding affinity and expression pattern. The binding affinity of each cAMP receptor is correlated with the stage of development during which it is expressed to optimally meet the needs of the cell at that time (Kim *et al*. 1996).

Members of the class C glutamate receptor-like protein family (which includes *grlG*) in *D. discoideum* also exemplify complexities of GPCR signaling*. GrlE*, which is the GABA receptor during late development, is also bound by the competitive inhibitor of GABA, glutamate. Depending on which ligand is bound, GABA or glutamate, *grlE* is coupled to a different G protein subunit (Gα7 or Gα9, respectively) (Anjard and Loomis 2006). Sporulation is delayed while glutamate is bound and then induced when displaced and bound by GABA, together ensuring the appropriate timing of sporulation (Anjard and Loomis 2006; Anjard *et al*. 2009). Another example of a glutamate receptor-like protein binding multiple ligands is the folic acid receptor *far1* (otherwise known as *grlL*), which also recognizes lipopolysaccharide (LPS) to mediate chemotaxis and phagocytosis of bacterial prey (Pan *et al*. 2016, 2018). And similar to the previously described multiple receptors of cAMP, other ligands bind multiple glutamate receptor-like proteins of class C as well. While *grlE* is the GABA receptor during late development, *grlB* is instead the GABA receptor during early stages of development (Wu and Janetopoulos 2013). There are also several examples in *D. discoideum* of potential redundancy (or rather, the suspected involvement of multiple receptors) of glutamate receptor-like proteins, wherein the loss of the focal receptor does not completely abolish the response as would be expected if a single receptor were responsible for ligand recognition and response. Examples in which the loss of the focal receptor does not completely abolish the response include the autocrine proliferation repressor protein A (*aprA*) receptor, *grlH* (Tang *et al*. 2018); the phenylthiourea receptor, *grlJ* (Robery *et al*. 2013); and the folic acid (and LPS) receptor, *far1* (*grlL*) (Pan *et al*. 2016, 2018).

Regarding the potential redundancy of *far1* (*grlL*), as previously mentioned, the involvement of *grlG* has been suggested (Pan *et al*. 2016; Lamrabet *et al*. 2020) but that has little relevance in this study as we have no reason to believe that our results are predation related. However, we do not rule out the possibility *grlG* may also interact with folic acid or another molecule with overlapping roles in bacterial predation as well as in cooperation and fruiting body formation. Given our understanding of the complex nature of GPCR signaling, even if *grlG* was found to be involved in the response to bacterial prey, that would not contradict the evidence we present here suggesting its role in cooperation. Instead, perhaps the knowledge that *grlG* and other members of the glutamate receptor-like protein family sometimes show low levels of overlapping functionality may help explain why the loss of *grlG* in our evolved clones did not always lead to the total loss of cooperation and non-fruiting.

## Conclusion

In this study we returned to cell lines previously evolved under low-relatedness by Kuzdzal-Fick *et al*. (2011) to investigate the genomic changes underlying the observed evolution of non-fruiting obligate cheaters. That study demonstrated how low relatedness in a natural population would lead to a collapse of multicellularity and with it, the advantages of fruiting body formation and spore dispersal. In this study we used whole-genome sequencing and variant analysis of those experimentally evolved *D. discoideum* low-relatedness cell lines. We identified parallel evolution of the orphan GPCR *grlG* as well as another 28 variants in other genes. The genes that we identified with variants are spread throughout the genome and do not share any significantly enriched functions or other annotations. In addition, none of these genes have been previously implicated in cheating or cooperation. The lack of overlap among these genes alone, nor with previously implicated cheater genes, is further testament to the complexity of the social cycle and aggregative multicellularity in *D. discoideum* and further highlights that these processes are orchestrated by a large diversity of genes and pathways.

The orphan GPCR *grlG* acquired variants of moderate to high impact in more than half of the experimental lines suggesting that the loss of *grlG* was adaptive under the experimental conditions of low-relatedness. By screening additional clones from the evolved lines, we identified a significant correlation between the complete loss of cooperation (inability to form the social stage, a fruiting body) and the presence of variants in the 5’ region of *grlG* containing the signal peptide and extracellular binding domain. Our results suggest that *grlG* is one of now two genes (second to *fbxA* (Ennis *et al*. 2000)) to have been implicated in obligate non-fruiting cheating in *D. discoideum*. Although future work is needed to identify the interacting molecules and the precise role of *grlG*, this work adds *grlG* to the growing list of GPCRs in the *D. discoideum* genome to have a role in multicellular development. This is noteworthy because it brings us one step closer to deorphanization of the ever-important GPCRs and highlights the need to expand research efforts to characterize the remaining GPCRs in *D. discoideum*.

## Acknowledgments

We thank members of the Queller-Strassmann laboratory for helpful advice on this research and Jennie Kuzdzal-Fick for making her evolved lines available. This material is based upon work supported by the National Science Foundation under grant numbers IOS 16-56756 and DEB 17-53743. Additional funding came from a pilot grant from the McDonnell Genome Institute.

## Supplemental Files

### Supplemental Methods

#### Supplemental Tables

**Additional File 1** Results of the limited screening of clones for variants in genes other than *grlG*

**Supplemental Material 1** Example images of non-fruiting clone morphology

**Supplemental Material 2** Variant calls that survived filtration from each of the three variant callers (GATK, Freebayes and Delly), in VCF format

**Supplemental Material 3** *Dictyostelium discoideum* gene annotations downloaded from NCBI on 10-25-19

**Supplemental Material 4** Tables showing the resulting number of variants after each step of filtration of the VCF files independently generated by each caller.

**Supplemental Material 5** Results of the functional annotation clustering with a high classification efficiency performed for the 29 genes with SNPs using the online tool DAVID (https://david.ncifcrf.gov/)

