## Supplemental Methods for "Loss of altruism in the social amoeba *Dictyostelium discoideum* is associated with the G protein-coupled receptor *grlG*"

### *Variant Read Support*

For the final confirmation of variant support and quality we generated read counts for each SNP across all samples using bam-readcount v0.7.4 (https://github.com/genome/bam-readcount). Because we expect that a true variant that is identified in a clone will be detected at some level in its population, we removed any SNP for which the origin population did not support the same alternate allele with at least five reads (as mentioned in the variant calling descriptions). We also confirmed for each SNP that it was not supported by reads in any other evolved lines or the ancestor, which could indicate poor sequence quality, shared ancestral polymorphism or contamination.

For the majority of the SNPs that passed filtration, only the evolved line from which the SNP was called contained reads supporting the alternate allele; there were a few exceptions which we removed from further analyses. We removed one variant in the gene, *dhkE,* which was only called in one line (23), but it was represented (albeit by only a small number of reads) in all of the evolved lines, with an average allele frequency of 6.3%. We also identified one evolved line (18) that contained an unusually large number of reads supporting the alternate allele for three SNPs that were only called in other (non-18) evolved lines. The extent of the allele support (27%, 31% and 17%) in 18 suggests that they are either true shared polymorphisms or evidence of contamination. Two of these sites (in DDB_G0276291 and DDB_G0276367) were called in line 17 and the other site (in DDB_G0276529) was called in lines 7 and 16 (this was the only SNP called in more than one population). In addition to those shared SNPs there was also a shared deletion (in *grlG*) that was called in lines 17 and 18. Each of these shared variants are highly supported and we believe them to be true positive calls. But we cannot rule out potential contamination of the population sample of line 18, so we reject the called variants from that sample. Excluding calls made in that sample does not impact any SNPs. The three SNPs described here were not called in line 18 nor were there any other SNPs called in that sample. The deletion that was called in lines 17 and 18 will not be removed but we only report the call made in line 17. The deletion in the population sample of line 17 has been PCR verified as part of the analysis described in the section, *Association of Variants with the Loss of Altruism.* No other SVs were called in the population sample for line 18.
