## Additional File 1 for "Loss of altruism in the social amoeba *Dictyostelium discoideum* is associated with the G protein-coupled receptor *grlG*"

### ***Variants in Other Genes (not grlG) are Less Likely to be Associated with the Loss of Cooperation***

For a subset of the clones that we screened for *grlG* variants we also screened for the presence of one or more SNPs that we called in other gene(s) (**Supplemental Table S4**). These additional screens were mostly uninformative, but we did identify one other apparent correlation between a variant and the clonal phenotype. We found a perfect correlation between a SNP called in DDB_G0278531 (a premature stop codon near the center of this uncharacterized membrane protein) and the fruiting phenotype in evolved line 24. We screened 24 clones (12 non-fruiters and 12 fruiters) from that line and found that all 12 non-fruiting clones carried the SNP while the 12 fruiting clones did not carry the SNP. This is precisely the pattern that we expect to see if a mutation is completely penetrant and is responsible for the loss of cooperation. However, all but one of the non-fruiting clones we screened that have the SNP in DDB_G0278531, also have a deletion in the 5' region of *grlG* making it difficult to know which variant is responsible. But unlike the parallel evolution seen in *grlG*, this is the only variant called in the gene and it was only called in the one evolved line.
