## Supplemental Material 1 for "Loss of altruism in the social amoeba *Dictyostelium discoideum* is associated with the G protein-coupled receptor *grlG*"

**Supplemental Material 1.** Example images of non-fruiting clone morphology. Clones displayed are (A) 19-NF1, (B) 14-NF1, (C) 20-NF1, (D) 15-NF1, (E) 7-NF3 and (F) 5-NF2. The NF clone IDs were retained for record keeping purposes; they are not related to the number of clones sequenced for a line.

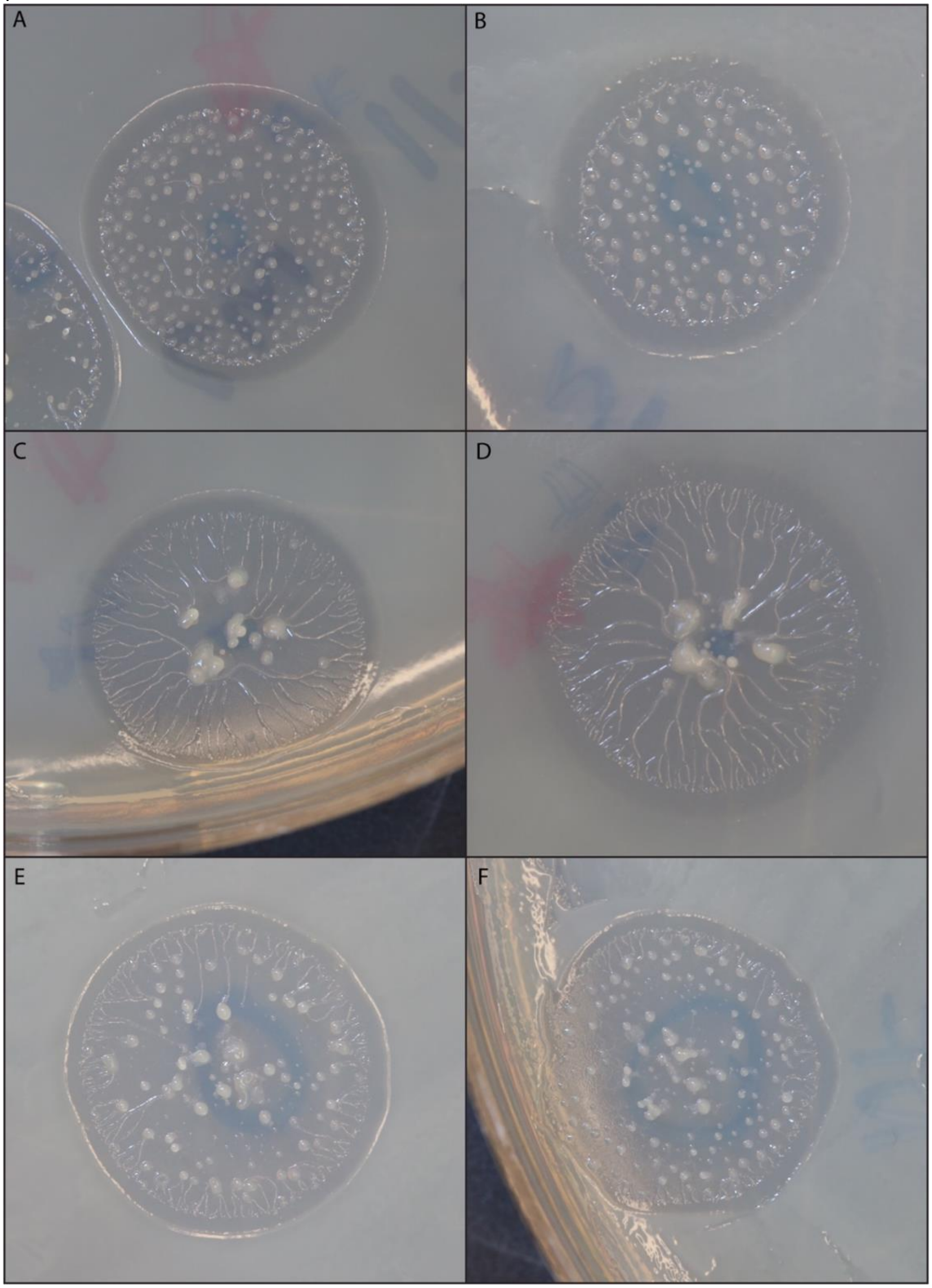
