## Supplemental Material 4 for "Loss of altruism in the social amoeba *Dictyostelium discoideum* is associated with the G protein-coupled receptor *grlG*"

**Supplemental Material 4.**
The resulting number of variants retained after each step of filtration of the VCF files independently generated with GATK (**A**), Freebayes (**B**) and Delly (**C**). The three resulting VCFs are available in Supplemental Material 2.

**A. GATK4 Variant Filtration**

| | **GATK joint-called variants** |
|---|---|
| Full VCF (SNPs, Indels, etc.) | 53,220 |
| SNPs only (no more than 2 alleles) | 36,794 |
| Major allele frequency (MAF) in the ancestor > 0.9 | 19,569 |
| Remove Ancestral Non-reference sites (background) | 2,029 |
| GATK Hard Filters (quality, strand bias, etc.) | 1,477 |
| Max missing 10 | 1,383 |
| Alternate allele (AC) called <= 3 samples | 746 |
| QUAL > 200 | 316 |
| Maximum depth < 1.5X average 4864 and 1914 | 306 |

**B. Freebayes Variant Filtration**

| | **Freebayes joint-called variants** |
|---|---|
| Full VCF (SNPs, Indels, etc.) | 32,755 |
| Normalization | 144,581 |
| Subset to simple SNPs only | 118,293 |
| Remove sites with MAF in Ancestor < 0.90 | 73,672 |
| Remove Ancestral Non-reference sites (background) | 55,573 |
| MQM 40 | 50,297 |
| Alleles <= 3 | 1,931 |
| Max missing 10 | 1,871 |
| QUAL > 200 | 1,570 |
| SAF>0 & SAR > 0 | 354 |
| SAP>0.5 & SRP> 0.5 | 266 |
| RPR>1& RPL>1 | 230 |

**C. Delly Variant Filtration**

| | **Delly joint-called variants** |
|---|---|
| Full VCF | 10,139 |
| Germline filter applied (Delly) | 131 |
| PASS filter (PE > 3 or PE > 5 for translocations, QUAL >= 20) | 29 |
| Alternate allele (AC) called < 3 samples | 16 |
| Complex variants discarded | 12 |
